# Impaired spatial memory in adult vitamin D deficient BALB/c mice is associated with reductions in spine density, nitric oxide, and neural nitric oxide synthase in the hippocampus

**DOI:** 10.1101/2022.01.12.476116

**Authors:** Md. Mamun Al-Amin, Robert K. P. Sullivan, Suzy. Alexander, David A. Carter, DanaKai. Bradford, Thomas H. J. Burne

**Affiliations:** Queensland Brain Institute, The University of Queensland, Brisbane 4072, Australia; Queensland Centre for Mental Health Research, Wacol 4076, Australia; Australian E-Health Research Centre, CSIRO, Pullenvale 4069, Australia

**Keywords:** Vitamin D, hippocampus, spatial learning, CA1, mushroom spine

## Abstract

Vitamin D deficiency is prevalent in adults and is associated with cognitive impairment. However, the mechanism by which adult vitamin D (AVD) deficiency affects cognitive function remains unclear. We examined spatial memory impairment in AVD-deficient BALB/c mice and its underlying mechanism by measuring spine density, long term potentiation (LTP), nitric oxide (NO), neuronal nitric oxide synthase (nNOS) and endothelial NOS (eNOS) in the hippocampus. Adult male BALB/c mice were fed a control or vitamin D deficient diet for 20 weeks. Spatial memory performance was measured using an active place avoidance (APA) task, where AVD-deficient mice had reduced latency entering the shock zone compared to controls. We characterised hippocampal spine morphology in the CA1 and dentate gyrus (DG) and made electrophysiological recordings in the hippocampus of behaviourally naïve mice to measure LTP. We next measured NO, as well as glutathione, lipid peroxidation and oxidation of protein products and quantified hippocampal immunoreactivity for nNOS and eNOS. Spine morphology analysis revealed a significant reduction in the number of mushroom spines in the CA1 dendrites but not in the DG. There was no effect of diet on LTP. However, hippocampal NO levels were depleted whereas other oxidation markers were unaltered by AVD deficiency. We also showed a reduced nNOS, but not eNOS, immunoreactivity. Finally, vitamin D supplementation for 10 weeks to AVD-deficient mice restored nNOS immunoreactivity to that seen in in control mice. Our results suggest that lower levels of NO, reduced nNOS immunostaining contribute to hippocampal-dependent spatial learning deficits in AVD-deficient mice.

## Introduction

Vitamin D deficiency is a global public health burden, affecting millions of people worldwide. Low serum vitamin D levels are associated with many neuropsychiatric diseases, such as schizophrenia [1, 2], autism [3, 4] and depression [5, 6]. In these diseases, hippocampal structural deformation has been reported as a central issue [7–9]. In recent years, there has been an increasing amount of literature suggesting that vitamin D deficiency is associated with a reduced hippocampal volume [10, 11] and disrupted hippocampal structural connectivity [12, 13].

The hippocampus is an important brain region contributing to spatial learning and memory, partly by consolidating short to long term memory. Vitamin D may have a specific role in the hippocampus, since neurons in the hippocampus and its various subfields express the vitamin D receptor (VDR) [14]. Moreover, vitamin D supplementation has been shown to improve hippocampal function [15]. Collectively, the data to date suggest vitamin D deficiency may play a role in hippocampal-dependent spatial learning [16]. However, the impact of adult vitamin D (AVD) deficiency on hippocampal-dependent spatial learning has been poorly understood with little agreement on the association of spatial learning deficits with AVD deficiency [17]. However, both AVD deficiency [15, 18], and developmental vitamin D (DVD) deficiency were associated with reduced spatial learning in rats [19]. To further investigate the effects of AVD deficiency on spatial learning and memory formation in mice, we used an active place avoidance (APA) task. The APA task is hippocampal-dependent, with performance based on the animal’s ability to encode and retrieve the spatial memory of a location [20, 21].

Structural transformations of the synapse have been demonstrated following learning of a spatial learning task, resulting in an increased number of dendritic spines in CA1 neurons compared to animals who fail to show learning [22–24]. Importantly, AVD deficiency was shown to reduce the hippocampal synaptic proteins, e.g., synaptojanin and synaptotagmin in the rat hippocampus [15]. Furthermore, spatial learning and hippocampal mushroom spine density were associated with the expression of synaptojanin [25]. This evidence indicates that vitamin D might have an impact on synapse formation, specifically through spine morphology. With respect to morphology, there are four types of dendritic spines - stubby, filopodia, thin and mushroom. The spines are highly dynamic and alter their shape over time [26]. However, repetitive training stabilizes the spine structure [27], and spatial learning performance was shown to increase the density of mushroom spines in hippocampal CA1 neurons [24]. Based on the existing evidence, we hypothesized that AVD deficiency would alter spine morphology in hippocampal CA1 pyramidal neurons.

A previous study in adult rats showed that eight weeks of a vitamin D depleted diet led to decreased induction of long term potentiation (LTP) in the hippocampus of anesthetized animals, compared to rats fed on a control diet or vitamin D supplemented diet [28]. LTP is important in learning and memory and it is well known that glutamate neurotransmission plays a central role in LTP [29].

AVD deficiency was not shown to alter proliferation or survival of new neurons in the hippocampus of BALB/c mice [30]. However, AVD deficiency resulted in an imbalance between excitatory (increased GABA) and inhibitory (depleted glutamate) neurotransmission in the whole brain of BALB/c mice [31]. Vitamin D has been shown to provide antioxidative properties [32]. The expression of glutathione in rat astrocyte cells was enhanced with vitamin D supplementation [33], which may be due to alterations in glucose-6-phosphate dehydrogenase enzyme which is required for glutathione synthesis [32]. Glutathione is one of the major antioxidant enzymes and the hippocampus is very susceptible to oxidative damage due to a high oxygen requirement. Therefore, we hypothesized that the oxidative stress markers such as, glutathione, nitric oxide (NO), lipid peroxidation and protein oxidation could be altered in AVD deficiency.

Neuronal nitric oxide synthase (nNOS) releases NO in the mammalian brain [34], and facilitates the release of neurotransmitters [35] and synaptic communication [36]. nNOS is found within the spine head [37], thereby regulating synaptic communication in the presynaptic and post-synaptic terminals for GABA [38] and glutamate [39]. Studies have shown that vitamin D deficiency downregulates NO synthesizing enzymes [40], and vitamin D supplementation enhances this enzyme [41]. Therefore, AVD deficiency may have an impact on GABAergic inhibitory neurons through neuronal nitric oxide synthase (nNOS) and/or neuropeptide y (NPY). Besides nNOS, the NPY-positive interneurons were also shown to regulate hippocampal excitatory transmission [42] and ameliorate spatial learning deficits [43]. Hippocampal nNOS cells are often colocalized with NPY interneurons [44]. Thus, nNOS and NPY positive interneurons are present in the hippocampus and potentially contribute to hippocampal networks [45, 46].

In this study, we first aimed to measure hippocampal-dependent spatial memory in BALB/c mice using the APA task. We characterised spine morphology in the dendrites of pyramidal neurons in the CA1 and DG regions of the hippocampus, and measured synaptic plasticity using electrophysiological recordings within the hippocampus of behaviorally naïve mice. We next measured the level of neurochemicals such as glutathione, malondialdehyde (lipid peroxidation), advanced oxidation of protein products (AOPP) and NO in hippocampal tissue. Finally, we quantified the labelling of hippocampal nNOS, eNOS and NPY in hippocampal subfields.

## Experimental procedures

### Animals and housing conditions

We purchased 61 male BALB/c mice (age 10 weeks) from Animal Resources Centre, Canning Vale, WA, Australia. The BALB/c mice were housed in groups of 4, in OptiMICE cages (Animal Care Systems, CO, USA), with corn cob bedding (Sanichips, Harlan Laboratories, USA) at the Animal Facility, Queensland Brain Institute, The University of Queensland, Australia. The animal housing conditions were maintained with a 12-h light-dark cycle. All of the animals had free access to water and food. The animals were given either a control diet (Standard AIN93G Rodent diet with 1500 IU vitamin D3/kg, Specialty Feeds, WA, Australia) or a vitamin D-deficient diet (Vitamin D Deficient AIN93G Rodent diet, Specialty Feeds, WA, Australia) (Supplementary table S1). Following behavioural testing, a separate group of AVD-deficient BALB/c mice (*n* = 6) were returned to the control diet (vitamin D containing diet) for 10 weeks. The experimental work was completed with approval from the University of Queensland Animal Ethics Committee (QBI/376/15/NHMRC), under the guidelines of the National Health and Medical Research Council of Australia.

### Active place avoidance test

We tested mice (*n* = 15/group) in an active place avoidance (APA) task using methods previously described [47–49]. The APA task has been used to assess hippocampal-dependent spatial learning and memory formation in rodents [49]. In this task, mice need to avoid a shock zone on the platform using four external cues hanging on the nearby walls. The apparatus is made by Bio-Signal Group, which consists of an elevated arena with a grid floor. A transparent circular boundary (77 cm diameter, 32 cm high, made of plexiglass) is used as a fence and placed on the elevated arena. The stage rotated clockwise (1 rpm) and delivered an electric shock through the grid floor.

The shock zone was defined within a 60° region of the stationary room and kept constant in relation to the room coordinates. The position of the mouse was tracked by PC-based software that analysed images from an overhead camera and delivered shocks appropriately (Tracker, Bio-Signal Group Corp., Brooklyn, NY). A mouse was placed opposite the shock zone facing the wall and trained to avoid an unmarked invisible shock zone using the four external cues. Entrance into the shock zone resulted in the delivery of a brief constant mild electrical foot shock (60 Hz at 0.5 mA for 500 ms). If the mouse remained in the shock zone, it received additional shocks of the same intensity at 1.5-second intervals until the animal moved out of the area. The experiment was conducted over five days. A duration of 5 min habituation session on Day 1 without shock was followed by 10 min testing session for four consecutive days. We collected data for the latency to enter into the shock zone, the number of shocks received and the distance travelled using Track Analysis software (Bio-Signal Group).

### Perfusion and brain tissue collection

Mice (*n* = 11/group) were anesthetized by intraperitoneal (i.p.) administration of phenobarbital injection (0.1 ml, 3.34 ml/kg body weight). Perfusion was carried out transcardially using 50 ml of ice-cold phosphate-buffered saline (PBS) at a pH of 7.6, 0.9% sodium chloride solution (50 ml) followed by fixative solution (0.01% formaldehyde) (50 ml), and preserved in PBS overnight in 4°C.

### Immunohistochemistry

We used a rotary microtome to slice 10 μm coronal sections and collected a one-in-ten series of the coronal sections to stain nNOS and NPY interneurons (*n* = 5/group). The sections were dried at 40°C on a hot plate for 24h, before being placed in Antigen Recovery Solution in an orbital shaker/incubator for 30 min at 40°C. Sections were blocked at room temperature for 1h with a blocking solution containing 3% normal goat serum, 0.05% saponin, 0.1% Triton X-100 and 10% sodium azide in PBS. Goat primary antibody of nNOS (at dilution of 1:100), rabbit primary antibody of NPY (at dilution of 1:1,000) and mouse primary antibody of eNOS (at dilution of 1:2,000) were purchased from Thermo Fisher Scientific. The antibodies were used to incubate the sections for 48h at room temperature. Secondary antibodies, Alexa fluor-555 anti-goat, Alexa fluor-555 anti-mouse and Alexa fluor-488 anti-rabbit fluorescent markers, were used at room temperature for 12 h. The sections were quickly washed with PBS twice followed by three washes with PBS at 15 min intervals. The sections were finally stained with the nuclei marker DAPI for 15 min (Sigma, 1:5,000). Vectashield (Vector laboratories, USA) mounting medium was used to mount the sections and they were stored at 4°C.

### Golgi-cox impregnation

Perfused brains (*n* = 6/group) were immersed in a Golgi-cox solution for 2-weeks. Golgi-cox staining was performed according to [50] with a few modifications. Briefly, 5% potassium dichromate was dissolved in double distilled water (DDW) to prepare solution-A. Similarly, 5% mercuric chloride and 5% potassium chromate were separately dissolved in DDW to prepare solution-B and -C under light protected glass container using a magnetic stirrer. Then 5 parts of solution-A and 5 parts of solution-B were added slowly with continuous stirring. Next, solution-C (4 volume) was diluted with 10 volume DDW (4: 10) and mixed slowly with the mixture of solution-A and -B to prepare working solution. This working solution was filtered to remove precipitates and kept in a light-protected container.

Brain sections (thickness 150 μm) dried onto gelatin coated slides at 4°C were cut using a vibratome (Lecia 1000s) in a 30% sucrose solution. The sections were briefly rinsed with DDW, then, incubated in 30% ammonia solution in the dark for 10 min. The sections were washed with DDW for 5 min. Subsequently, 1% sodium thiosulfate was added and sections incubated in the dark for 10 min. The sections were washed in DDW then dehydrated using graded ethanol and xylene. Mounting was performed using DPX (Dibutylphthalate Polystyrene Xylene) under the fume hood.

### Microscopy

The fluorescence images were captured using a Zeiss Axio Imager Z1 microscope. Cells that immunolabelled with nNOS and NPY were included in the imaging as a tiled image and stitched subsequently. An experimenter blind to the treatment conditions counted the number of nNOS-positive cells, NPY-positive cells and co-localized nNOS and NPY cells. A custom pipeline developed in Cellprofiler [51] was used for semi-automatic counting. The area of the hippocampal subfield was measured in ImageJ (Fiji). Four anatomically matched dorsal sections (starting 1.34 - 2.46 mm posterior to bregma) were included in the counting protocols. Spine images were taken using a microscope (Axio Imager, Carl Zeiss) with a 100x oil objective. The following criteria were considered while selecting the spine of neurons; i) branch should be standalone, clearly identifiable from the neighbouring neuron, ii) not truncated iii) spines were clearly visible. Images of 4/5 branches of a neuron were taken by a person blind to the treatment conditions. Spine images were captured on spines present on processes at least 10 μm long at the apical dendritic branch of CA1 area. We also captured dendritic spines present in the molecular layer of the upper and lower dentate gyrus. Z-stack images were captured with 0.12 μm slice interval. We images 100-120 dendrites per sample (*n* = 6/group). We analysed a total of approximately 1280 dendrites from 12 animals, by selecting 4-5 dendrites per neuron; 3 neurons per hemisphere and 4 sections per sample. There were 50/60 dendrites per sample (total sample =12). We selected 4/5 dendrites per neuron, 3 neurons per hemisphere and 4 sections per sample.

### Spine analysis

The dendrites were manually traced and the dendritic spines were traced using a point-and-click method within neurolucida-360 in neurolucida software [52]. Neurolucida explorer software was used to manually edit the spines. The spine density was analysed by the number of spines present in the 10 μm length of the dendritic processes. The spine classification was performed in neurolucida-360 (MBF Bioscience). Neurolucida categorized spine into three classes; mushroom, stubby and thin. Neurolucida software does not have any feature to classify filopodia, which may be due to the structural similarity between filopodia and thin spines.

### Electrophysiology

Mice (n=4 Control and n=5 AVD-deficient) were anaesthetized and transcardially perfused with ice cold cutting solution containing (in mM) 118 NaCl, 2.5 KCl, 25 NaHCO_3_, 1.2 NaH_2_PO_4_, 10 D-glucose, 0.5 CaCl_2_, 3 MgCl, (pH 7.4, osmolarity = 300 - 310 mmol/Kg) saturated with 95% O_2_/5% CO_2_. Brains were rapidly removed, glued to a stage and 400 μm coronal whole-brain slices were prepared while submerged in cold cutting solution using a vibrotome VT1000S (Leica, Wetzlar, DEU). Slices were recovered for 40 min at 32 °C in saturated ACSF (cutting solution with 2.5 mM CaCl_2_ and 1.3 mM MgCl) before being returned to room temperature for 2-7 hours. Slices were transferred to a submerged recording chamber constantly perfused with saturated ACSF and maintained at 32 °C.

Extracellular recordings were made using glass microelectrodes (5 – 7 MΩ) filled with ACSF from stratum radiatum (CA1 region) or stratum moleculare (dentate gyrus). Stimulating micropipettes (approx. 1 MΩ) were placed either anterograde or retrograde of the recording electrode on the surface of the slice and input was recruited by an isolated constant voltage (100 μs duration, 0.05 Hz, AMPI, Jerusalem, ISR). To investigate input recruitment to CA1 or DG, stimulus intensity was increased from 0 to 50 V, in 10 V increments, and the slope of the average of three field excitatory post-synaptic potentials (fEPSPs) measured. All subsequent protocols in each slice utilised stimulus intensities to elicit a fEPSP 30% of maximum amplitude. Paired pulse ratio was determined by varying the interval between two stimuli to 20, 50, 150 and 400 ms. Long-term synaptic plasticity was examined by applying three trains of high-frequency stimulation (HFS, 100 Hz for 1 s) separated by five min intervals. Input was followed at least 60 min after first conditioning stimulus. Recordings in CA1 were discarded if fEPSP slope deviated > ± 25% from the start of the baseline period. In four experiments, a second stimulating pipette was placed and interleaved opposite the main stimulus electrode, but no conditioning stimulus was applied. We measured the voltage dependence of Schaffer collateral-mediated post-synaptic currents (PSC) after application with antagonists (NBQX, AP5, and PTX) for AMPA, NMDA or GABA_A_ receptors.

Output signal (20 kHz) was recorded using an Axopatch 700A (Molecular Devices, Sunnyvale, CA, USA) with a CV-7B head-stage (Molecular Devices, Sunnyvale, CA, USA). Recordings were sampled at 10 kHz and digitized via an ITC-16 (Instrutech Corp., Greatneck, NY, USA) connected to Axograph X software (Axograph Scientific, www.axograph.com). The slope and amplitude of an average of five or more fEPSP were measured using the linear curve or peak fitting in Axograph X software, respectively. An 8-pole Bessel filter (2.5 kHz cut-off) was applied to representative traces.

### Determination of Nitric oxide (NO)

Mice (*n* = 6/group) were anesthetized by intraperitoneal (i.p.) administration of phenobarbital injection (0.1 ml, 3.34 ml/kg body weight) and the hippocampus dissected on ice and snap frozen in liquid nitrogen. Nitric oxide was assayed according to a previous method [53, 54] using Griess reagent system [55] with a few modifications. In this experiment we used 0.1% w/v NED solution (naphthyl ethylene diamine di-hydrochloride) instead of 1-napthylamine (5%). The reaction mixture containing hippocampal tissue homogenate (50 μl supernatant) and phosphate-buffered saline (50 μl) were incubated at 25°C for 15 min. Then 50 μl of sulphanilamide solution (1% sulphanilamide in 5% phosphoric acid) was added and allowed to sit for 5 min. The absorbance was measured at a wavelength of 540 nm against the corresponding blank solutions. Sodium nitrite was used as a standard sample.

### Determination of reduced Glutathione (GSH)

Glutathione in the hippocampal tissue was measured based on a previous method [56]. Hippocampal tissue homogenate was prepared using 0.6% sulfosalicylic acid and Triton-X solution, centrifuged at 8000g for 10 min at 2–4 °C and the clear supernatant transferred to a fresh tube for total GSH assay. The assay mixture contained 20 μl of supernatant, 20 μl of KPE (potassium phosphate-EDTA buffer) and 120 μl of DTNB [5,5-dithio-bis (2-nitrobenzoic acid)], and GR (glutathione reductase) solutions. The reaction was initiated with the addition of 60 μl β-NADPH (Nicotinamide adenine dinucleotide 2’-phosphate). The absorbance was taken instantly at 412 nm UV-spectrophotometer every 40s for 3.2 min (5 readings).

### Determination of oxidized Glutathione (GSSG)

GSSG in the hippocampal tissue was measured based on a previous method [56]. Hippocampal tissue homogenate of sulfosalicylic acid extract (100 μl) was mixed with 2 μl of 2-vinylpyridine and left for 1 h at room temperature under a fume hood. Then 6 μl of triethanolamine was added vigorously and allowed to sit for 10 min for the neutralization process. The reaction was initiated with the addition of 60 μl β-NADPH. The absorbance was taken instantly at 412 nm UV-spectrophotometer every 40s for 3.2 min (5 readings).

### Determination of advanced oxidation of protein products (AOPP)

We followed spectrophotometric procedure to determine the AOPPs as previously described [57, 58]. Briefly, 50 μl of supernatant was diluted (1:2) with PBS. Next, 100 μl of 1.16 M potassium iodide and 50 μl of acetic acid were mixed to read the absorbance at 340 nm wavelength. The concentration of AOPP was expressed in μmol/mg of tissue. PBS was used as blank and chloramine T (0–100 μmol/l) were used to prepare the calibration curve.

### Determination of Malondialdehyde (MDA)

We measured lipid peroxidation colorimetrically according to a previous protocol [59, 60]. Briefly, 100 μl of tissue homogenate was treated with 100 μl of TBA-TCA-HCl (1:1:1 ratio) reagent (2-thiobarbituric acid 0.37%, 0.25 N HCl and 15% TCA). The solution was placed in a water bath at 70°C for 15 min. The solution was then cooled and the clear supernatant was separated to read the absorbance at 535 nm wavelength. Then 1, 1, 3, 3-tetramethoxypropane was used to prepare standard curve. The level of MDA was expressed as nmol/mg of tissue.

### Determination of serum 25-(OH)D

At the end of the behavioural experiment we collected a blood sample from the saphenous vein of one mouse per cage (*n* = 6/group) to confirm vitamin D levels. A liquid chromatography-tandem mass spectrometer (Sciex Instruments, ON, Canada) on a 4000 QTrap API AB mass spectrometer was used to measure the levels of 25(OH)D in serum samples [61]. As expected, there was a significant (*t*_10_ = 3.96; *p* < 0.01) difference on the level of serum 25(OH)D with AVD-deficient mice having lower serum 25(OH)D (M = 4.87, SEM = 0.29) compared to the controls (M = 23.97, SEM = 4.81).

### Data analysis and statistics

Data were analysed with SPSS (Version 25.0) software. We used independent sampling (1 data point/animal) and tested the data for normality and equal variances. We used parametric statistical tests and conducted repeated measure ANOVA (considering the variables Day and Diet) to analyse the data obtained from the active place avoidance test. Unpaired t-test was also conducted to analyse the effects of Diet (control or AVD-deficient) on the levels of nitric oxide, glutathione, lipid peroxidation, protein oxidation, nNOS, NPY and eNOS labelling. The *p*-values less than 0.05 were considered significant. Data are presented as mean±SEM.

## Results

### AVD deficiency impaired spatial learning in the active place avoidance test

To test for spatial learning and memory, we trained BALB/c mice in the APA test. We found a significant main effect of Day (*F_3,108_* = 8.00;*p* < 0.001) and Diet (*F_1,108_* = 5.18;*p* = 0.024) on the latency to enter the shock zone (Fig. 1A). The AVD-deficient mice had a lower latency to enter the shock zone than control mice. Multiple comparison showed that the control mice had a significantly higher latency on Day 4 compared to AVD-deficient mice. We also conducted a partial ANOVA to test the improvement of learning throughout the experiment in the control and AVD-deficient mice. There was a significant main effect of Control Diet (*F_1,21_* = 7.56; *p* = 0.005) but not AVD-deficient Diet (*F_1,21_* = 1.85; *p* = 0.181) on the latency to enter the shock zone, suggesting that the AVD-deficient mice had impaired learning of the APA task.

**Fig. 1.**
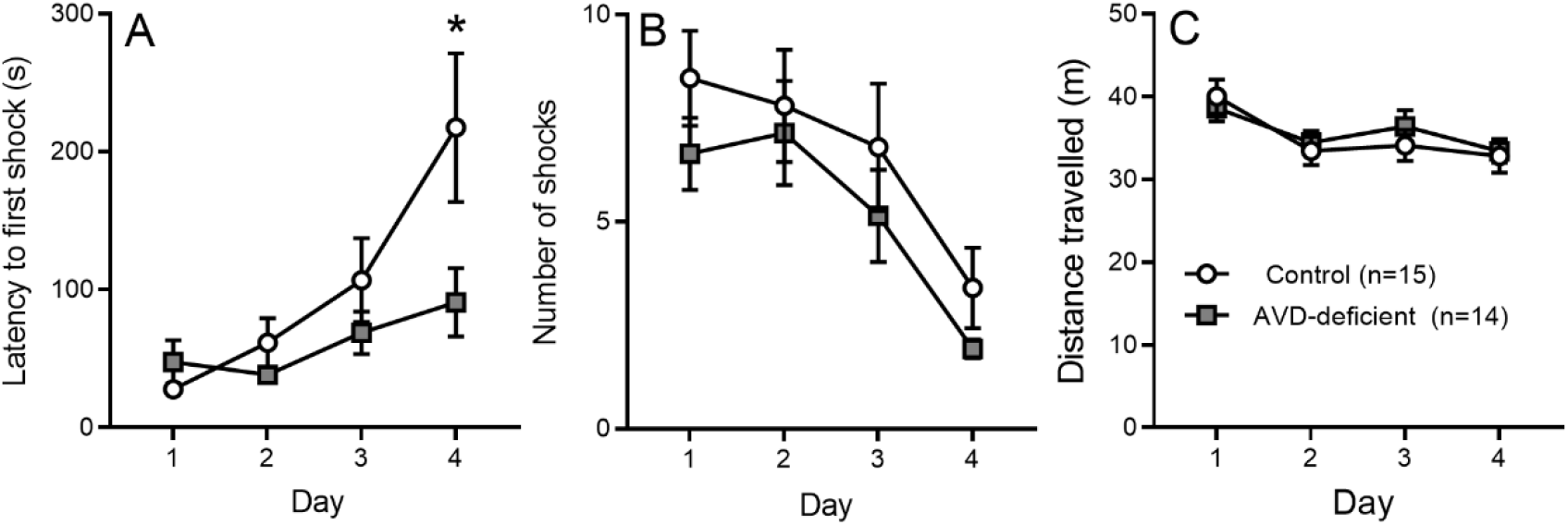
The AVD-deficient BALB/c mice showed reduced spatial learning in the active place avoidance (APA) task. AVD-deficient mice had a lower latency to enter the shock zone (A). The number of shocks (B) and distance travelled (C) over 4 days of the testing sessions were not varied between the two diet groups. The groups were: control and AVD-deficient. Values are mean ± *SEM; n* = 14/15 per group; *\*p* < 0.05.

There was a significant main effect of Day (*F_3,108_* = 8.10;*p* < 0.001), but not Diet (*F_1,108_* = 3.04; *p* = 0.08) on the total number of shocks (Fig. 1B). Further partial ANOVA test showed that the control and AVD-deficient mice showed a significant main effect on the number of shocks, suggesting that both groups of mice were able to reduce their exposure to shocks with repeated training. We also observed a significant main effect of Day (*F_3,108_* = 4.72; *p* < 0.01), but not Diet (*F_1,108_* = 0.24; *p* = 0.624) on the total distance travelled throughout the experimental sessions (Fig. 1C). So while AVD-deficient mice entered the shock zone earlier, they did not differ from controls in terms of the number of shocks received nor running ability in the APA task.

### AVD deficiency altered spine morphology in CA1

To understand the underlying basis of spatial learning deficits (Fig. 2A), we measured the hippocampal dendritic spine morphology using a Golgi-cox staining technique (Fig. 2B). We observed no significant main effect of Diet on total spine density in the DG and CA1 subfields (Fig. 2C-E). We then analysed the main effect of Diet on the density of each spine type e.g., thin, stubby and mushroom. We found a significant main effect of Diet (*F_1,11_* =14.2; *p* = 0.004) on the mushroom spine density in the CA1 proximal dendrites (Fig. 2E) but not in the DG upper or lower blade (Fig. 2C,D). The AVD-deficient mice had a reduced number of mushroom spines compared to the controls. No differences were seen in the other spine types.

**Fig. 2.**
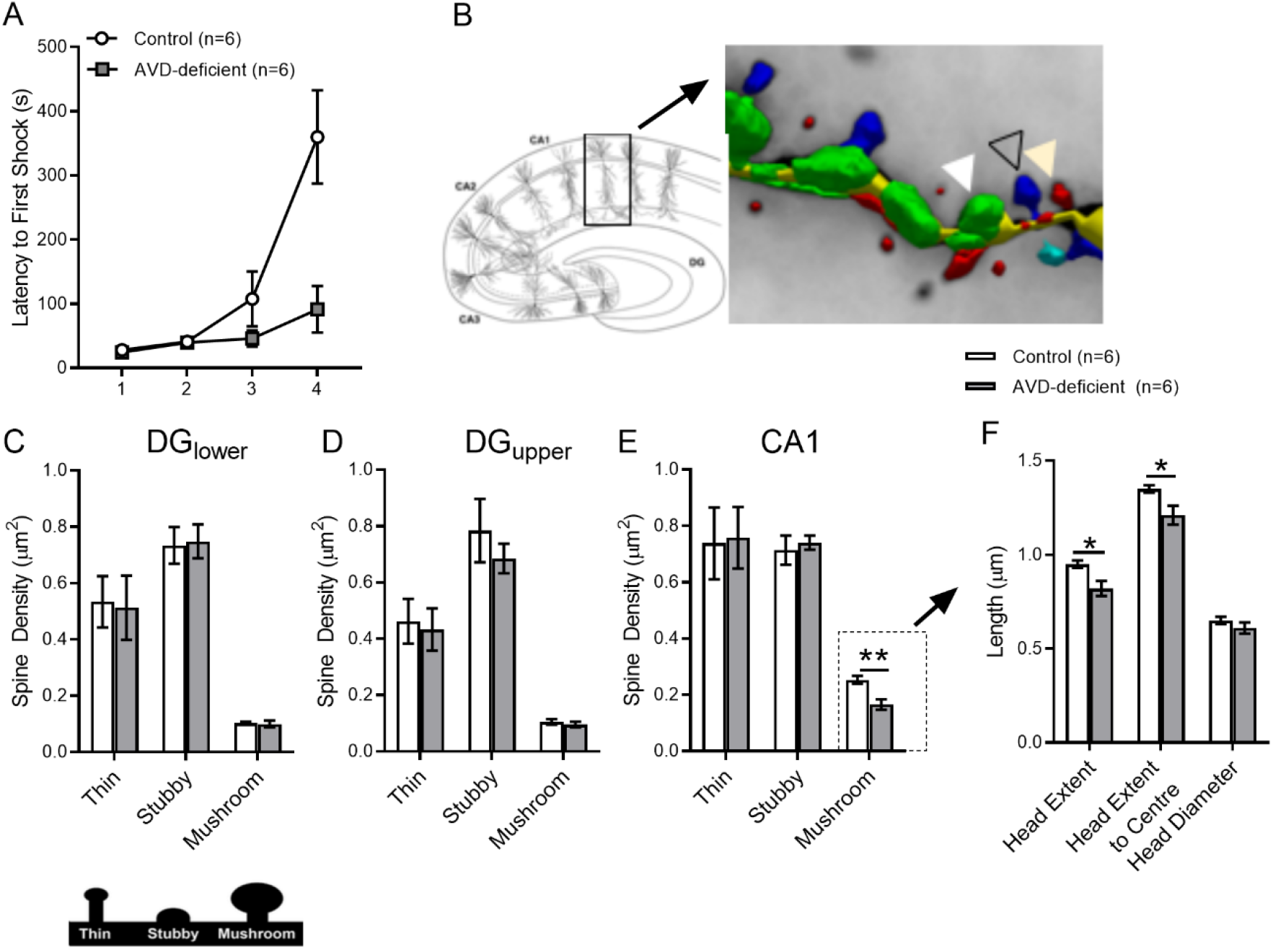
The AVD-deficient mice that were included in the Golgi-cox experiment showed a lower latency to enter the first shock (A). The hippocampal CA1 pyramidal neuron (B) and spine types are shown where, the white, black and yellow arrowheads represents stubby (green), thin (red) and mushroom (blue) spines respectively. AVD deficiency reduced the density of CA1 hippocampal mushroom spines in the BALB/c mice. There was no effect of Diet in the dentate gyrus (DG) lower (C) or upper (D) blade, but a significant main effect of Diet on the spine type in CA1 (E). The head morphology, including head extent and the head extent to centre of the mushroom spine (F), was reduced in AVD-deficiency. The groups were: control and AVD-deficient. Values are mean ± *SEM; n* = 6 per group; *\*p* < 0.05.

Further analysis on the morphology of mushroom spines showed a significant main effect of Diet on the Head Extent (*F_1,11_* = 10.0; *p* = 0.01) and Head Extent to centre (*F_1,11_* = 7.5; *p* = 0.02) of the mushroom spines (Fig. 2F). AVD-deficient mice had a reduced head extent and a decreased head extent to the centre of the mushroom spines.

### AVD deficiency did not impair LTP in CA1 pyramidal neurons

To address the role of AVD deficiency on synaptic plasticity, we have studied LTP in CA1 pyramidal neurons using high-frequency stimulation (HFS) of the Schaffer collaterals paired with depolarisation. We could induce LTP to neurons in both control and AVD-deficient mice to a similar extent (Fig. 3). Assessment of the voltage dependence of Schaffer collateral-mediated post-synaptic currents (PSC) using antagonists (NBQX, AP5, and PTX) indicated no effect of diet on AMPA, NMDA or GABA_A_ receptors, respectively (data not shown).

**Fig. 3.**
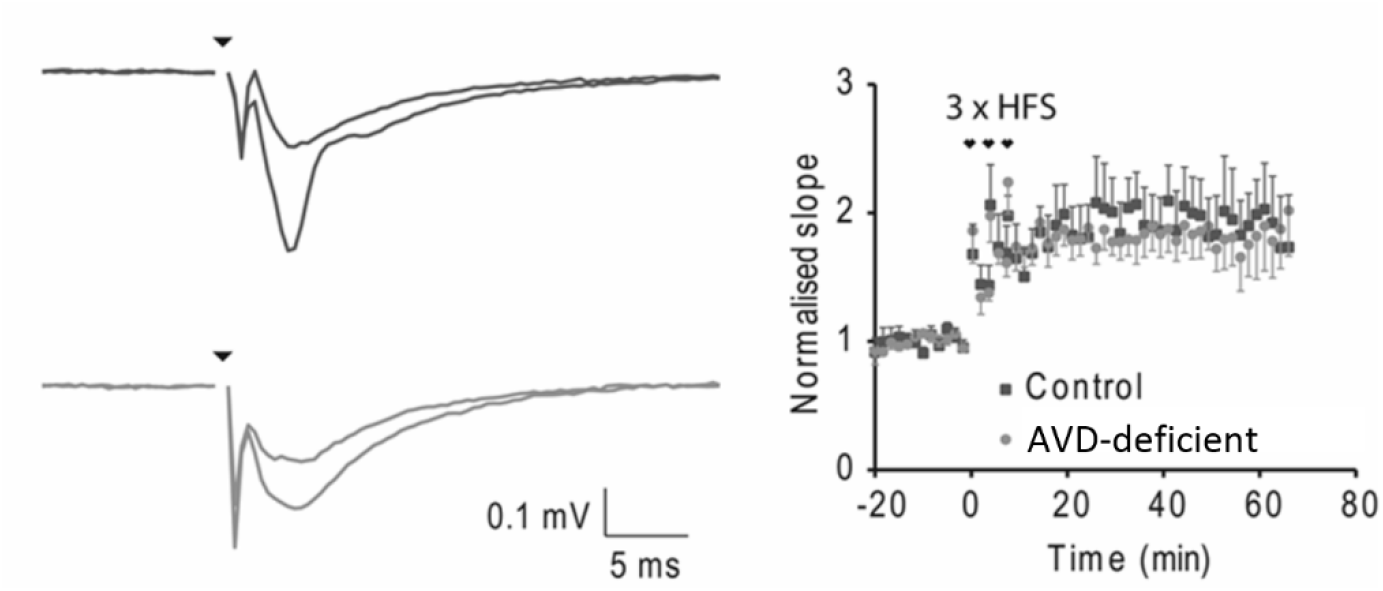
AVD deficiency did not alter LTP in CA1 pyramidal neurons. Using high-frequency stimulation (HFS) of the Schaffer collaterals paired with depolarisation we could induce LTP to neurons in both control and AVD-deficient mice to a similar extent. n = 4/5 per group

### AVD deficiency was associated with reduced NO levels in the hippocampus

There was no significant difference in the level of total glutathione (GSH + GSSG) (*t_10_* = 0.55; *p* = 0.591, GSH (*t_10_* = 0.39; *p* = 0.703), GSSG (*t_10_* = 1.22; *p* = 0.247) (Fig. 4) and GSH to GSSG ratio (*t_10_* = 1.38; *p* = 0.196) (Fig. 4). However, there was a significant effect of Diet (*t_10_* = 2.64; *p* = 0.024) on the level of NO in the hippocampus (Fig. 4). In addition, there was no significant effect of Diet on MDA (*t_10_* = 0.92; *p* = 0.378) (Fig. 4) or AOPP (*t_10_* = 0.15; *p* = 0.5876) (Fig. 4) levels in the hippocampal tissue.

**Fig. 4.**
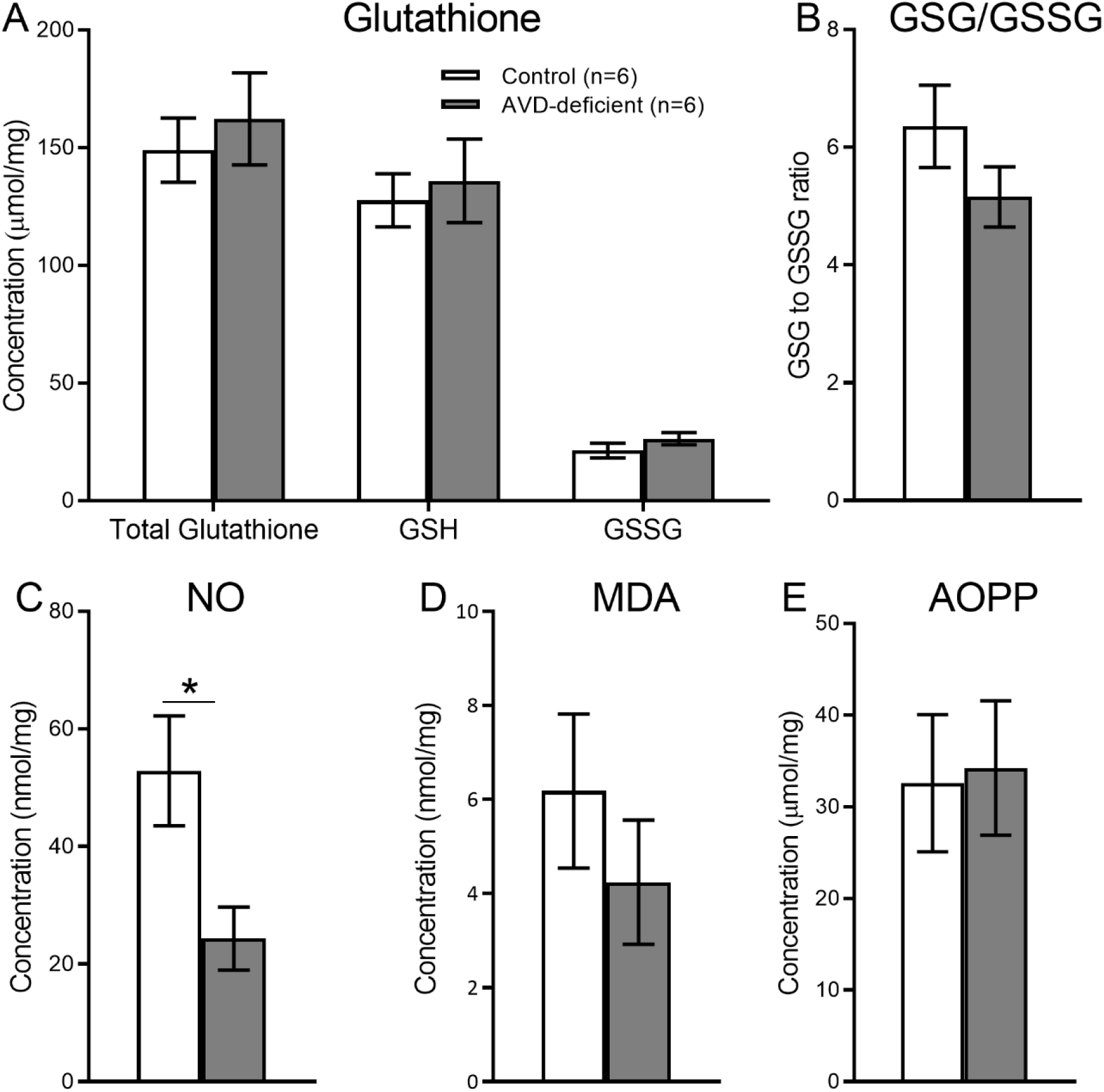
AVD deficiency depleted hippocampal nitric oxide in BALB/c mice. The level of NO was depleted in the hippocampal tissue in AVD deficiency (C). However, there was no significant main effect of Diet on the level of glutathione oxidized (GSSG) (A), glutathione reduced (GSH), total glutathione (GSH+GSSG), glutathione ratio (GSH/GSSG) (B), malondialdehyde (MDA) (D), advanced oxidation of protein products (AOPP) (E) in the hippocampal tissue. The groups were: control and AVD-deficient mice. Values are mean ± SEM; n = 6 per group; *p < 0.05.

### AVD deficiency was associated with reduced hippocampal nNOS expression

An independent samples *t*-test showed a significant difference between control and AVD-deficient mice on the number of nNOS^+^ cells in the CA1 (*t_8_* = 3.76;*p* = 0.018), CA2 (*t_8_* = 6.38;*p* < 0.001), CA3 (*t_8_* = 3.46; *p* = 0.011) and DG (*t_8_* = 3.17; *p* = 0.019; representative immunostaining is shown in Fig 5A-H). The number of nNOS^+^NPY^−^ cells were significantly lower in the AVD-deficient hippocampal subfields (Fig. 6). Moreover, AVD-deficient mice had significantly fewer nNOS^+^NPY^+^ cells in the CA1 (*t_8_* = 2.79; *p* = 0.026) and CA2 (*t_8_* = 3.70; *p* = 0.015) but not in the CA3 (*t_8_* = 2.30; *p* = 0.074) or DG (*t_8_* = 1.03; *p* = 0.343). The nNOS^−^NPY^+^ cells were not significantly affected in any of the hippocampal subfields by AVD deficiency; CA1 (*t_8_* = 2.01; *p* = 0.101), CA2 (*t_8_* = 0.92; *p* = 0.388), CA3 (*t_8_* = 0.618; *p* = 0.074) and DG (*t_8_* = 0.68; *p*=0.515). We also measured fluorescent intensity of nNOS immunoreactivity in the hippocampal subfields. *There was a* significant difference between control and AVD-deficient mice on the percentage change of fluorescent intensity of nNOS cells in the hippocampus (*F_1,8_* = 20.73, *p* = 0.002, repeated measures ANOVA), and this difference was seen in each subregion; CA1 (*t_8_* = 3.83;*p* = 0.005), CA2 (*t_8_* = 6.40;*p* < 0.001), CA3 (*t_8_* = 3.69; *p* = 0.006) and DG (*t_8_* = 2.41; *p* = 0.04).

**Fig. 5.**
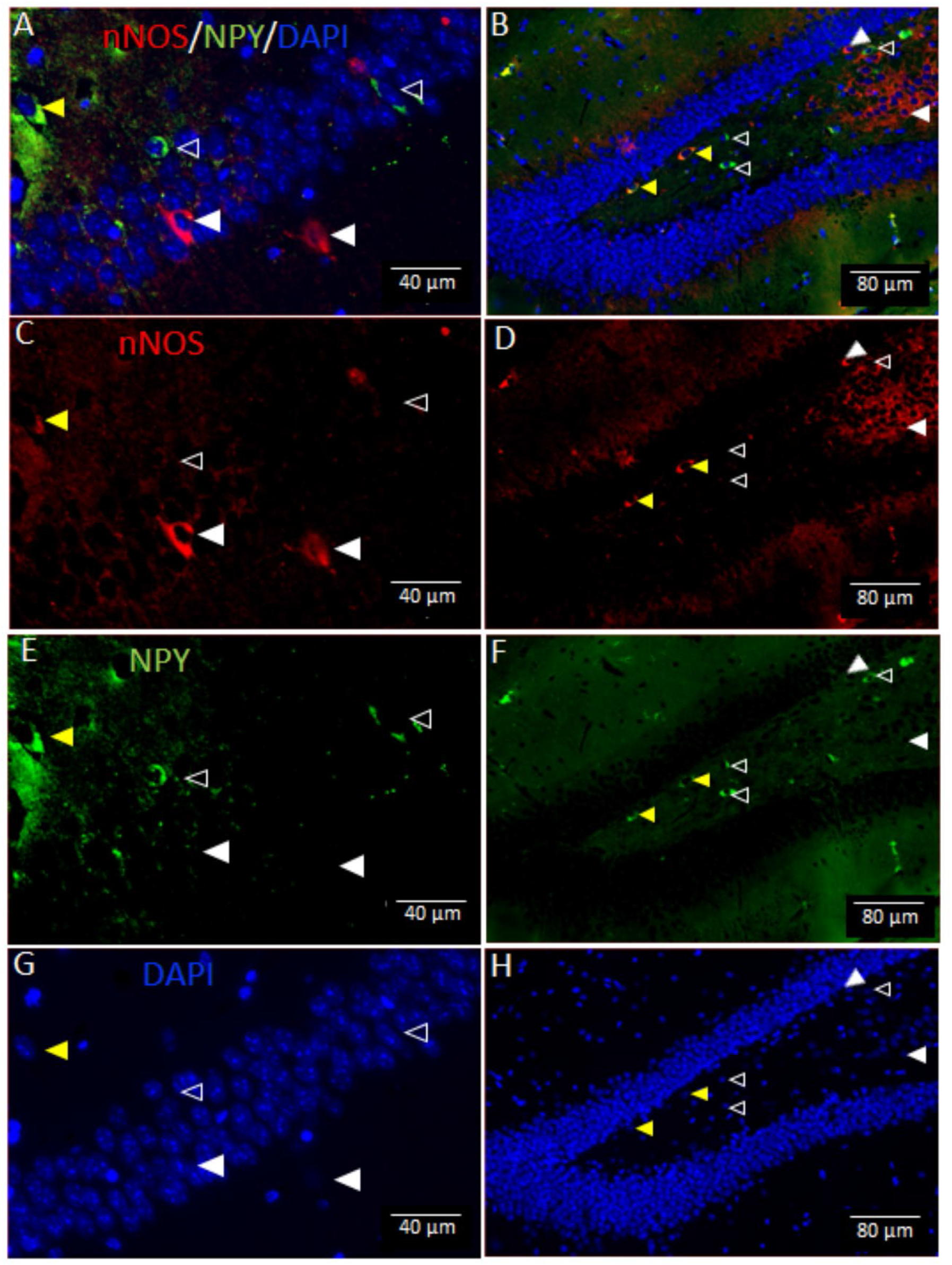
Vitamin D deficiency reduced the expression of hippocampal neuronal nitric oxide synthase (nNOS) in BALB/c mice. The top panels show triple labelling (A,B) of nNOS (C,D), neuropeptide-Y (NPY) (E,F) and DAPI (G,H) in the hippocampus. Arrowheads indicate nNOS^+^NPY^−^ (white), nNOS^+^NPY^+^ (yellow) and nNOS^−^NPY^+^ (transparent) cells.

**Fig. 6.**
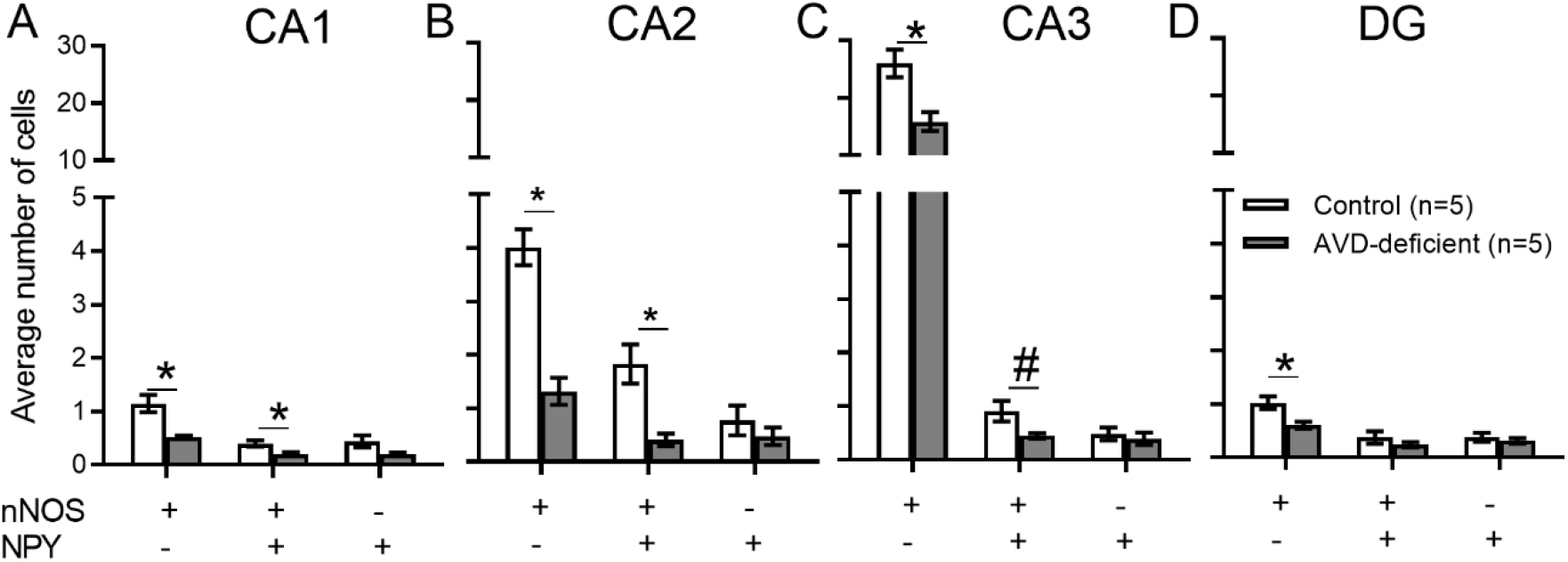
Vitamin D deficiency reduced the expression of hippocampal neuronal nitric oxide synthase (nNOS) in BALB/c mice. The vertical axis represents the average number of cells and the horizontal axis represents various types of cells (nNOS or NPY) in each sub region of the hippocampus; CA1 (A), CA2 (B), CA3 (C) or DG (D). Values were mean ± *SEM*; *n* = 5 per group; \**p* < 0.05; *\*\*p* < 0.01; *#p* < 0.06.

### AVD deficiency was not associated with hippocampal eNOS immunoreactivity

An independent samples *t*-test did not show any significant difference between control and AVD-deficient mice on the number of eNOS^+^ cells in the CA1 (*t_6_* = 0.22; *p* = 0.83), CA2 (*t_6_* = 0.45; *p* = 0.67), CA3 (*t_6_* = 0.62; *p* = 0.0.56) and DG (*t_6_* = 0.06; *p* = 0.96) (Fig. 7).

**Fig. 7.**
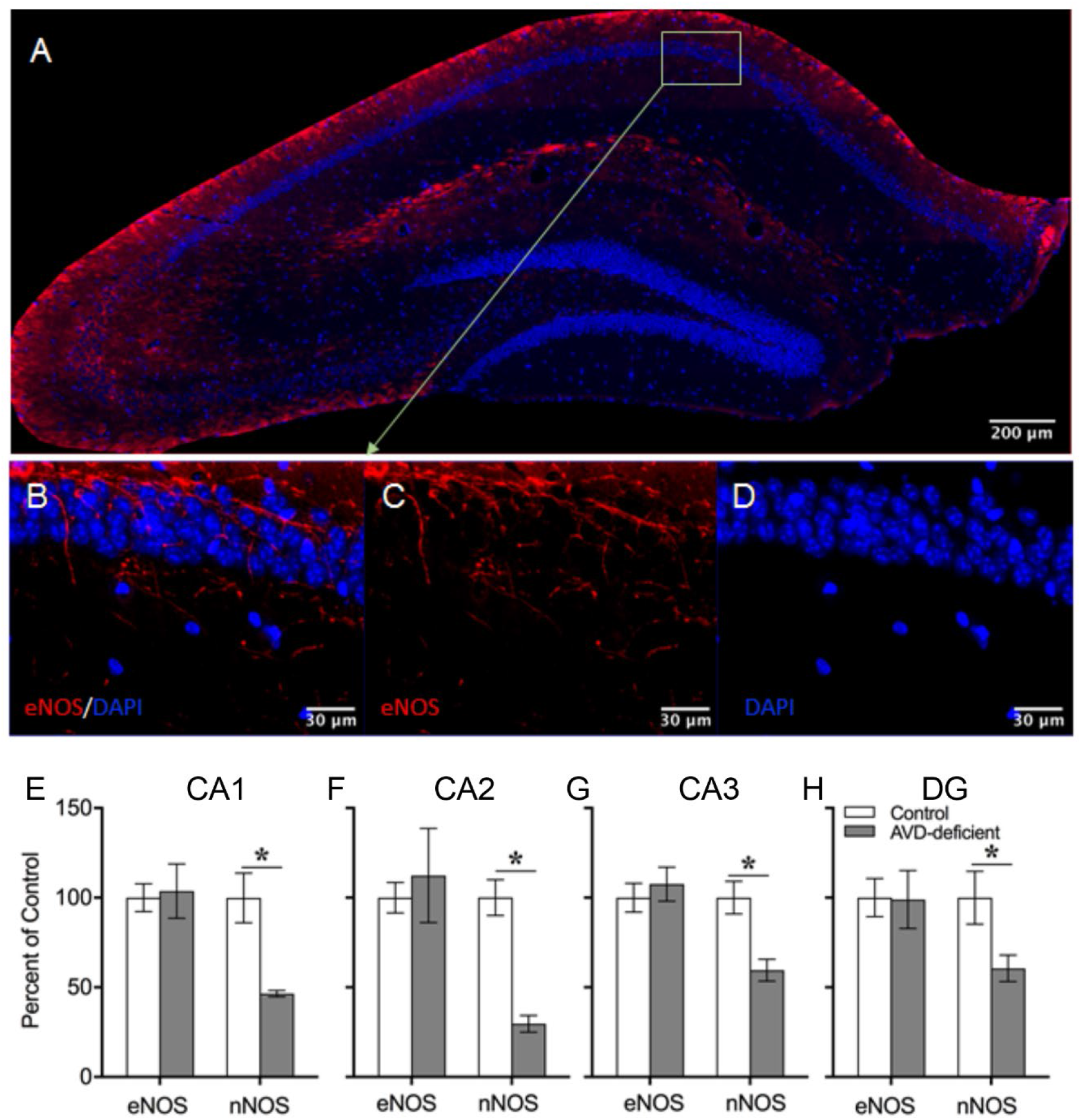
Adult vitamin D deficiency did not alter eNOS immunoreactivity in the hippocampus of BALB/c mice. There was no significant difference in eNOS immunoreactivity in mouse hippocampal subfields following 10-weeks of adult vitamin D deficiency. The top panel shows double labelling of eNOS and DAPI of a coronal hippocampal section (A). The middle panel shows labelling of eNOS and DAPI of hippocampal CA1 region (B) eNOS (C) and DAPI (D). The bottom panel (E-H) showed measures of percentage fluorescent intensity of eNOS and nNOS in control and AVD-supplemented mice. Values are mean ± SEM, *n* = 5 per group. \**p* < 0.05.

### Chronic supplementation of vitamin D prevented the decline of nNOS immunoreactivity in AVD-deficient mice

#### Immunoreactivity of nNOS

There were no significant differences in the average number of nNOS counts between control and AVD-supplemented mice in CA1 (*t_6_* = 1.82; *p* = 0.118), CA2 (*t_6_* = 0.23; *p* = 0.823), CA3 (*t_6_* = 1.46; *p* = 0.192) and DG (*t_6_* = 0.411; *p* = 0.695) (Fig. 8).

**Fig. 8.**
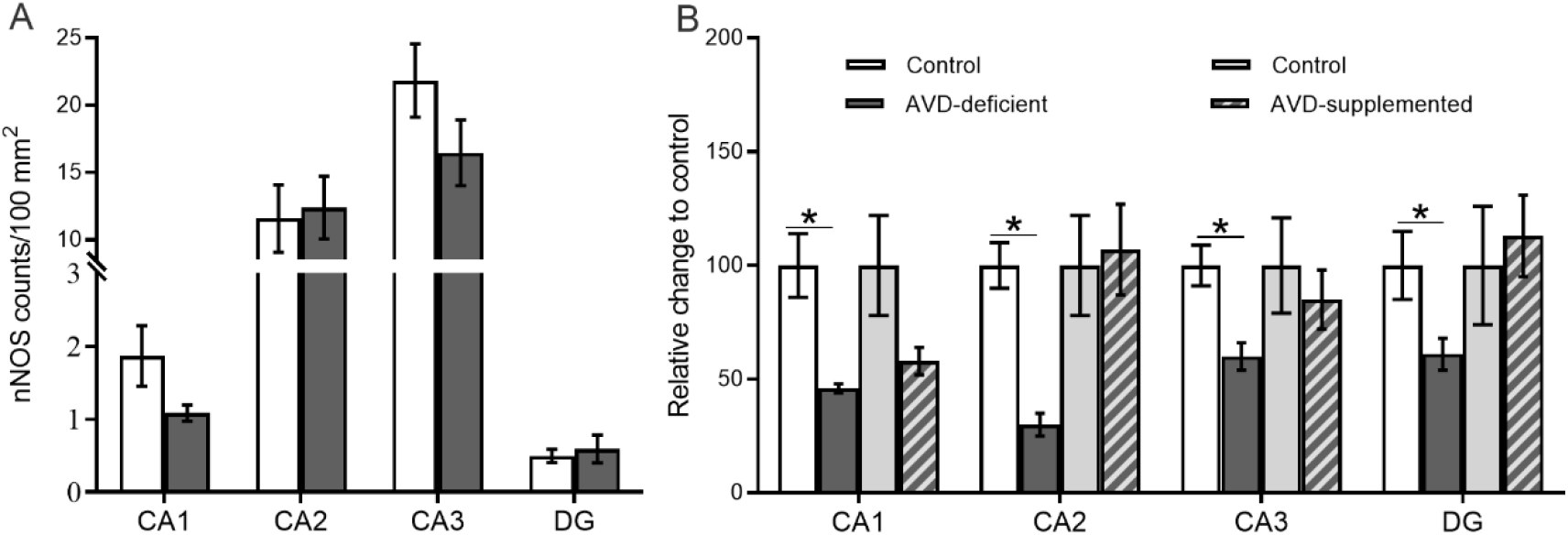
Chronic vitamin D supplementation prevents the reduction of nNOS immunoreactive cells in the hippocampus of AVD-deficient mice. There was no significant difference in nNOS immunoreactivity in mouse hippocampal subfields following vitamin D supplementation. nNOS counts/100 mm2 in control and AVD-supplemented mice (A). Percentage change of nNOS counts in AVD-deficient mice relative to control was significantly lower in hippocampal subfields. However, relative change to control did not significantly vary in AVD-supplemented mice in hippocampal subfields (B). All mice were on a diet for 20 weeks. Values are mean ± SEM, *n* = 4/5 per group. \**p* < 0.05.

#### Relative change in nNOS immunoreactivity compared to control mice

The percentage change of nNOS counts in AVD-deficient mice relative to controls were significantly lower in CA1 (*t_8_* = 3.81; *p* = 0.01), CA2 (*t_8_* = 6.39; *p* = 0.007), CA3 (*t_8_* = 3.68; *p* = 0.008) and DG (*t_8_* = 2.39; *p* = 0.05) (Fig. 8). However, there were no significant differences in the percentage change of nNOS counts in AVD-supplemented mice relative to controls in the CA1 (*t_6_* = 1.81; *p* = 0.15), CA2 (*t_6_* = −0.23; *p* = 0.82), CA3 (*t_6_* = 0.61; *p* = 0.56) and DG (*t_6_* = −0.40; *p* = 0.69) (Fig. 7D).

## Discussion

This study investigated impaired spatial memory in AVD-deficient mice and examined potential neural correlates. There were four main findings from the present study. At the behavioural level, AVD-deficient mice had a shorter latency to enter the shock zone in the APA test, indicating a deficit in hippocampal-dependent spatial learning. We found that the density and morphology of mushroom spines in the hippocampal CA1 dendrites was diminished by AVD deficiency, despite no effect on LTP recorded in hippocampal slices. Potential cellular mechanisms underpinning this impairment included a depletion in the level of hippocampal NO in AVD deficient mice, despite no changes in other oxidative stress markers. Immunostaining of the hippocampal tissue showed a reduced labelling of nNOS but not NPY. These results suggest that the effect of AVD deficiency on spatial memory may be due to depletion of hippocampal NO levels, diminished nNOS immunoreactivity and reduced mushroom spine formation on CA1 pyramidal neurons in the hippocampus. A summary of the results are given in Table 1.

**Table 1.** Summary of the main results.

|  | GS<br>H | MD<br>A | AOP<br>P | NO | nNO<br>S+ | nNOS+<br>NPY- | nNOS+<br>NPY+ | NPY+ | nNOS-<br>NPY+ | eNOS | Mush<br>room | Stu<br>bby | Thi<br>n |
| --- | --- | --- | --- | --- | --- | --- | --- | --- | --- | --- | --- | --- | --- |
| CA1 | – | – | – | ↓ | ↓ | ↓ | ↓ | – | – | – | ↓ | – | – |
| CA2 |  |  |  |  | ↓ | ↓ | ↓ | ↓ | – | – | NA | NA | NA |
| CA3 |  |  |  |  | ↓ | ↓ | – | – | – | – | NA | NA | NA |
| DG |  |  |  |  | – | ↓ | – | – | – | – | – | – | – |

Nitric oxide (NO), Neuronal nitric oxide synthase (nNOS); Endothelial nitric oxide synthase (eNOS); neuropeptide-Y (NPY); Glutathione (GSH); Malondialdehyde (MDA); Advanced oxidation of protein products (AOPP); Cornu Ammonis (CA); Dentate Gyrus (DG); Decreased (↓); Not applicable (NA); Unchanged (–).

To understand the impact of AVD deficiency on hippocampal-dependent function, we tested BALB/c mice on a spatial learning task. The AVD-deficient mice had a shorter latency to enter the shock zone, indicating a deficit in spatial memory formation. In addition, the AVD-deficient mice did not show any significant improvement in their latency to enter the shock zone throughout the 4 days of training suggesting a reduced ability to learn the task. On the contrary, control mice showed improved spatial learning and memory formation following repeated training. This study did not find a difference between control and AVD deficiency on the number of shocks animals received during the experimental sessions. The parameter “distance travelled” throughout the experimental sessions also did not show any difference between the groups. This result clearly indicates that the AVD-deficient mice had similar motor activity. The APA task requires the mouse to retrieve the spatial information that was formed on the previous day, and once the mouse enters the shock zone they encode the spatial information of the invisible shock zone using distal visual cues. Therefore, we argue that the spatial learning performance in the APA test was purely hippocampal-dependent [21]. These results support the hypothesis that AVD deficiency impairs hippocampal-dependent spatial learning and memory formation [15, 18].

To understand the underlying basis of impaired hippocampal-dependent spatial learning, we measured spine morphology from the CA1 apical dendrites and DG dendrites. The hippocampal-dependent spatial learning task was shown to promote spine formation [22] and spine density [62] by the alteration of spine distribution and morphological alterations in the CA1 pyramidal neuron [63]. In addition, spatial learning tasks were also shown to increase the spine density in the dentate gyrus (DG) [64]. We observed that AVD deficiency reduced the densities of the mushroom spine in the CA1 but not in the DG regions. Our finding is consistent with previous studies showing increased mushroom spine density in spatially trained rodents [24, 65]. By contrast, mushroom spines were not affected following training in the spatial learning task in the DG regions by AVD deficiency. One possible explanation is that the CA1 hippocampal synapses are selectively vulnerable to vitamin D deficiency and the maturation of spines is differentially regulated in the DG. Another explanation is that the effect of the spatial learning task on the DG was transient [66, 67]. It is also possible that the DG has a different compensatory mechanism to vitamin D deficiency, such as altered neurogenesis. However, hippocampal CA1 synapses may produce persistent changes following training in the APA task [68]. Therefore, performance in the APA task may have a selective and consistent impact on the CA1 region but not in the DG, which may be regulated by vitamin D.

We further analysed a detailed structure of the mushroom spine heads such as “head extension”, “head extension to centre” and “head diameter”. We showed a reduced extent of head of the mushroom spines indicating that the synaptic surfaces were reduced by AVD deficiency. The spine head is an important parameter for synaptic communication; the bigger the head, the larger the synaptic contact, resulting in greater synaptic strength. Previous studies showed that calcium ions regulate the head extension of spine [69, 70]. Since vitamin D regulates calcium homeostasis it is possible that vitamin D deficiency reduced head extension of the mushroom spine by regulating calcium ions.

Long term potentiation (LTP) is an important molecular correlate with memory formation, but it was not impaired in AVD-deficient compared to control mice. There are a variety of mechanisms responsible for the short and long-term persistence of LTP after a conditioning stimulus. These include post-translational mechanisms, such as the trafficking of AMPAR to the post-synaptic density, or transcription- and translation-mediated changes [71, 72]. However, AVD deficiency did not affect the release properties of the perforant pathway or Schaffer collaterals, or the ability of these inputs to recruit a post-synaptic depolarisation. Even a strong conditioning stimulus was insufficient to differentiate STP between diets. These data indicate that AVD deficiency induced changes to the density of mushroom spines are either not sufficient to effect baseline synaptic neurotransmission or are compensated by other mechanisms. Another possible mechanism for LTP inhibition is a shifting towards GABA neurotransmission in the hippocampus, as demonstrated by the enhancement of LTP with the inclusion of picrotoxin in some studies [73]. More work is required to determine the mechanism by which AVD deficiency may impact on LTP.

We then measured oxidative stress markers in hippocampal tissue to understand the molecular basis of the spatial learning deficit. Oxidative stress was shown to impair hippocampal-dependent spatial learning [74]. Except for NO, we did not show any impact of AVD deficiency on oxidative stress markers. Our novel result of reduced NO levels is in agreement with data showing that vitamin D may regulate NO release in the hippocampus [41]. Previous studies have shown that vitamin D regulates NO release by the activation of endothelial nitric oxide synthase [40] and inducible nitric oxide synthase [41, 75] enzymes.

NO could have a divergent role in the hippocampus including spatial memory formation [76–78] and facilitation of LTP [79]. Spatial learning using the radial arm maze was shown to increase the release of NO in the hippocampus [80]. In addition, in active avoidance learning, NO was associated with improved spatial learning via glutamate production in the hippocampus [81]. However, LTP is normal in mice with a targeted mutation in either nNOS or eNOS and LTP in CA1 may be NO independent [82].

Nitric oxide is synthesized from three nitric oxide synthase (NOS) enzymes; neuronal (nNOS), endothelial (eNOS) and inducible (iNOS). The activity of nNOS and eNOS are calcium-dependent, whereas the activity of iNOS is calcium-independent [83]. The eNOS principally expresses in endothelial cells [84], iNOS expresses in macrophages and glial cells in the pathological state [85], whereas nNOS is primarily expressed in GABAergic interneurons [86]. In addition, the expression of hippocampal nNOS was shown to increase following spatial learning [87]. We therefore, first focus on the hippocampal nNOS interneurons for four important reasons. First, the majority of NOS activity in the hippocampus is from nNOS acitivity (98%), and <2% is eNOS activity [82]. Second, nNOS is calcium-dependent, and the calcium homeostasis is regulated by serum vitamin D. Third, nNOS is expressed in GABAergic interneurons and we have previously shown altered levels of GABA neurotransmitter in the whole brain of AVD-deficient BALB/c mice [31]. Fourth, the interaction between the nNOS and PSD-95 (post-synaptic density) are essential for synapse formation [88, 89]. We found a reduced nNOS expression in the hippocampal subfields. We did not find any previous evidence showing the effect of vitamin D on nNOS expression. However, 1,25(OH)2D3 was shown to increase eNOS via a direct transcriptional regulation [40]. Therefore, the genomic action of vitamin D may contribute to the reduction of nNOS, which may be the underlying basis of spatial memory deficits.

Furthermore, eNOS may also be altered by AVD deficiency due to its similarity with nNOS with regards to calcium-dependency. Therefore, we compared the nNOS and eNOS immunoreactivity in the hippocampal subfields between AVD-deficient and control mice. Although, nNOS immunoreactivity was reduced in AVD-deficiency, eNOS immunoreactivity in the hippocampal subfields were unchanged. Our eNOS finding is not consistent with a previous study that showed a regulatory role of vitamin D on eNOS in mouse artery [40]. It is possible that vitamin D has a differential role in blood vessels and hippocampus. Further studies are required to measure the eNOS immunoreactivity within the blood vessel in the hippocampus. In our immunostaining experiment, eNOS immunoreactivity could be due to the presence of eNOS in the blood vessels, astrocytes [90] or in the dendritic spines [91].

A large number of nNOS interneurons are shown to co-express NPY [44, 92, 93]. A reduced nNOS expression could be due to the lower levels of NPY [94]. However, we found that the NPY^+^ cells were only reduced when colocalized with nNOS^+^ cells, otherwise the expression of NPY^+^ cells were not affected by AVD deficiency. We also measured nNOS immunoreactivity in AVD-deficient mice supplemented with vitamin D for 10 weeks. We did not show any difference in immunoreactivity of nNOS between control and AVD-supplemented mice. These data show that chronic vitamin D supplementation improves nNOS expressing components in the hippocampus suggesting a direct role of AVD deficiency on hippocampal nNOS expression and this should be investigated in future studies.

Taken together, these data suggest that the reduced levels of NO and immunoreactivity of nNOS may be associated with reduced CA1 mushroom spine density. For example, NO was shown to contribute to synaptic function, such as the facilitation of synapse formation and growth of synaptic terminals [95, 96], and enhancement of synaptic efficiency by LTP induction [37]. Moreover, NO and nNOS may be responsible for mushroom spine reduction via its role in LTP in CA1 neurons [97, 98]. Finally, the interaction between NO and nNOS signalling facilitates synapse formation [99, 100]. In summary, the reduction of hippocampal nNOS, NO and mushroom spines may affect hippocampal-dependent spatial learning impairment in AVD deficiency. We have also shown that vitamin D supplementation can rescue nNOS immunoreactivity within the hippocampus. Future studies should examine the extent of these changes and whether, for example, L-arginine or vitamin D supplementation are able to rescue the spatial learning deficits in AVD deficiency.

## Acknowledgements

This research was supported by the National Health and Medical Research Council grant APP1070081 to TB and a University of Queensland International PhD Scholarship to MA.

