## Supplementary information for "Impaired spatial memory in adult vitamin D deficient BALB/c mice is associated with reductions in spine density, nitric oxide, and neural nitric oxide synthase in the hippocampus"

***Supplementary Table S1*** Composition of Speciality feed Diet (SF09-088 AIN93G Rodent Diet)

Supplementary Table S1.1 Calculated Nutritional Parameters

| **Nutrient** | **Amount** |
| --- | --- |
| Protein | 19.40% |
| Total Fat | 7.00% |
| Crude Fibre | 4.70% |
| AD Fibre | 4.70% |
| Digestible Energy | 16.1 MJ/Kg |
| % Total calculated digestible energy from lipids | 15.90% |
| % Total calculated digestible energy from protein | 21.10% |

Supplementary Table S1.2 Base components

| **Name of the Ingredients** | **Rate of inclusion** |
| --- | --- |
| Casein (Acid) | 200 g/Kg |
| Sucrose | 100 g/Kg |
| Soya Bean Oil | 70 g/Kg |
| Cellulose | 50 g/Kg |
| Maize Starch | 404 g/Kg |
| Dextrinised Starch | 132 g/Kg |
| DL Methionine | 3.0 g/Kg |
| Calcium Carbonate | 13.1 g/Kg |
| Sodium Chloride | 2.6 g/Kg |
| AIN93 Trace Minerals | 1.4 g/Kg |
| Potassium Citrate | 2.5 g/Kg |
| Potassium Dihydrogen Phosphate | 6.9 g/Kg |
| Potassium Sulphate | 1.6 g/Kg |
| Choline Chloride (75%) | 4.1 g/Kg |
| Oxicap E2 | 0.14 g/Kg |
| AIN93 Vitamins | 15 g/Kg |
| Vitamin K 0.23% | 0.87 g/Kg |

Supplementary Table S1.3 Calculated total vitamins

| **Name of the Vitamin** | **Rate of inclusion** |
| --- | --- |
| Vitamin A (Retinol) | 6000 IU/Kg |
| Vitamin D (Cholecalciferol)# | None added (Deficient)  1500 IU/Kg (Control) |
| Vitamin E (a Tocopherol acetate) | 115 mg/Kg |
| Vitamin K (Menadione) | 3.5 mg/Kg |
| Vitamin C (Ascorbic acid) | None added |
| Vitamin B1 (Thiamine) | 9.1 mg/Kg |
| Vitamin B2 (Riboflavin) | 9.3 mg/Kg |
| Niacin (Nicotinic acid) | 45 mg/Kg |
| Vitamin B6 (Pyridoxine) | 11 mg/Kg |
| Pantothenic acid | 24.5 mg/Kg |
| Biotin | 300 μg/Kg |
| Folic acid | 3 mg/Kg |
| Vitamin B12 (Cyanocobalamin) | 152 μg/Kg |
| Choline | 2380 mg/Kg |
| #Vitamin D was not given to the Adult vitamin D deficient mice for 10 weeks. However, control mice received 1500 IU/Kg for the same duration. | |

Supplementary Table S1.4 Calculated Amino Acids

| **Name of the Amino acids** | **Rate of inclusion** |
| --- | --- |
| Valine | 1.26% |
| Leucine | 1.80% |
| Isoleucine | 0.87% |
| Threonine | 0.79% |
| Methionine | 0.84% |
| Cystine | 0.05% |
| Lysine | 1.49% |
| Phenylalanine | 0.99% |
| Tyrosine | 1.01% |
| Tryptophan | 0.27% |
| Histidine | 0.60% |

Supplementary Table S1.5 Calculated Total Minerals

| **Name of the Minerals** | **Rate of inclusion** |
| --- | --- |
| Calcium | 0.47% |
| Phosphorus | 0.35% |
| Magnesium | 0.08% |
| Sodium | 0.15% |
| Chloride | 0.16% |
| Potassium | 0.40% |
| Sulphur | 0.23% |
| Iron | 68 mg/Kg |
| Copper | 7.0 mg/Kg |
| Iodine | 0.2 mg/Kg |
| Manganese | 19 mg/Kg |
| Zinc | 46 mg/Kg |
| Molybdenum | 0.15 mg/Kg |
| Selenium | 0.3 mg/Kg |
| Chromium | 1.0 mg /Kg |
| Fluoride | 1.0 mg/Kg |
| Lithium | 0.1 mg/Kg |
| Boron | 2.5 mg/Kg |
| Nickel | 0.5 mg/Kg |
| Vanadium | 0.1 mg/Kg |

Supplementary Table S1.6 Calculated Fatty Acid Composition

| **Name of the Minerals** | **Rate of inclusion** |
| --- | --- |
| Myristic Acid 14:0 | Trace |
| Palmitic Acid 16:0 | 0.72% |
| Stearic Acid 18:0 | 0.27% |
| Palmitoleic Acid 16:1 | 0.01% |
| Oleic Acid 18: | 1.60% |
| Gadoleic Acid 20:1 | 0.01% |
| Linoleic Acid 18:2 n6 | 3.57% |
| a Linolenic Acid 18:3 n3 | 0.48% |
| Total n3 | 0.48% |
| Total n6 | 3.57% |
| Total Mono Unsaturated Fats | 1.62% |
| Total Polyunsaturated Fats | 4.05% |
| Total Saturated Fats | 0.99% |
